# Identification of new resistance QTL regions in loquat cultivar ‘Champagne’ against *Pseudomonas syringae* pv. *eriobotryae* group C

**DOI:** 10.1101/2024.03.04.583427

**Authors:** Shogo Koga, Ryusei Kawaguchi, Tsunami Tanaka, Shigeki Moriya, Naofumi Hiehata, Koji Kabashima, Atushi J. Nagano, Yukio Nagano, Shinji Fukuda

**Author notes:** Corresponding author: Saga University, Faculty of Agriculture, Saga, 840-8502, Japan.

## Abstract

Loquat canker, caused by *Pseudomonas syringae* pv. *eriobotryae*, is a bacterial disease that infects loquat (*Eriobotrya japonica*) and has been reported in several countries. Three pathotypes, A, B, and C, have been reported in Japan. The loquat cultivar ‘Champagne’ is resistant to the loquat canker group C and possesses a qualitative trait governed by a recessive homozygous *pse-c* gene located on Linkage Group 3 (LG3), and quantitative traits located at unidentified loci. In this study, we identified novel quantitative trait loci (QTL) regions for resistance to group C in this cultivar. A seedling population with ‘Tanaka’ (*Pse-c*/*Pse-c*) crossed with ‘Champagne’ (*pse-c*/*pse-c*) was tested. The genetic map of ‘Champagne’ includes a total of 1,016 SNP markers mapped across 17 LGs, covering a total distance of 1,301 cM and an average marker density of 1.4 cM/locus. In addition to minor potential QTLs, the major QTL for resistance to loquat canker group C was detected in the upper region of LG14, with the QTL contributing 6.9% to the disease index. The results of this study open new possibilities for resistance breeding against this disease.

**Highlights.:** A total of 1,016 SNP markers were mapped on a linkage map consisting of 17 linkage groups with a total distance of 1,301 cM.

QTL analysis revealed a novel resistance QTL region in ‘Champagne’ against loquat canker group C in the upper part of the LG14.

The identified QTLs in this study provide new possibilities for resistance breeding in loquat.

## 1. Introduction

The loquat (*Eriobotrya japonica* (Thunb.) Lindl.) is an evergreen fruit tree belonging to the subtribe Malinae, tribe Maleae, subfamily Amygdaloideae of the family Rosaceae (Campbell et al., 2007; Potter et al., 2007; Liu et al., 2020). It is believed that its cultivation began in southern China (Ding et al., 1995; Wang et al., 2017). Indeed, the cultivars introduced from China to Japan have led to the various Japanese cultivars seen today (Ichinose, 1995), a fact supported by DNA analysis (Nagano et al., 2022). Cultivated loquat was reportedly introduced from China to European countries, then from Europe to Florida between 1867 and 1870, and from Japan to California (Morton, 1987). However, although no records remain, DNA analysis has suggested the possibility that cultivars genetically distinct from the current Asian cultivars, not favored by Asians but instead preferred by Westerners, were introduced to the West (Nagano et al., 2022). Thus, this species has been introduced to many countries worldwide and is cultivated in more than 30 countries (Lin et al., 1999). The major producing countries are China, followed by Spain, Pakistan, and Japan (Lin, 2007).

In commercial loquat cultivation, the tree often suffers from bacterial diseases. Among these, loquat canker caused by *Pseudomonas syringae* pv. *eriobotryae* is a problematic disease that has been confirmed in many countries (Alippi and Alippi, 1990; Lai et al., 1971; Lin et al., 1999; McRae and Hale, 1986; Wimalajeewa et al., 1978). The disease affects all parts of the loquat, including buds, leaves, fruit, trunks, and underground parts (Morita et al., 1988; Mukoo, 1952; Suga et al., 2007). When the disease develops on the main trunk, it becomes a major issue in loquat cultivation because it leads to a decline in vigor and eventually a decrease in yield (Morita, 1978). Furthermore, the major Japanese cultivars ‘Mogi’, ‘Tanaka’, and ‘Nagasakiwase’ are susceptible to this disease (Morita, 1980; 1988), making complete control difficult. Therefore, the development of resistant cultivars against this disease is expected, which is promoting research on resistance genes.

Bacterial isolates from loquat production areas in Japan were classified into three groups: A (no pigment production and no leaf pathogenicity), B (no pigment production but with leaf pathogenicity), and C (pigment production but no leaf pathogenicity), based on the presence or absence of pigment production on the medium and leaf pathogenicity (halo formation) (Morita, 1978). Phylogenetic analysis using genome sequences showed that groups A and C are genetically similar, whereas group B is slightly different from them (Tashiro et al., 2021).

Resistance to group A is controlled by a gene named *Pse-a* (Hiehata et al., 2002). This gene locus is located at the upper end of the linkage group 10 (LG10) in loquats (Fukuda et al., 2014). Moreover, many cultivars resistant to group A also show resistance to group B, suggesting that the resistance mechanism against group B may originate from the *Pse-a* gene (Hiehata et al., 2002). In contrast, the Japanese cultivar ‘Shiromogi’ exhibits a recessive form of resistance to group C, governed by a different gene, *pse-c* (Hiehata et al., 2012), with this resistance locus located in the LG3 of loquats (Fukuda et al., 2019). Both the dominant *Pse-a* and the recessive *pse-c* contribute to strong resistance against the pathogen, and the trait characterized by *Pse-a* or *pse-c* is a qualitative trait, namely, a Mendelian trait.

Resistance to the pathogen might not be limited to the *Pse-a* and *pse-c* genes. The discovery of additional genes could significantly enhance resistance breeding efforts. Genetic resources showing resistance to group C are rare (Hiehata et al., 2014). Nonetheless, resistance to group C has been confirmed in a few varieties, including the American cultivar ‘Champagne’ in addition to ‘Shiromogi’. ‘Champagne’, which was bred in California around 1908 from unknown hybrid parents (Morton, 1987), was introduced to Japan in 1952. It possesses the resistance gene *Pse-a* against group A (Hiehata et al., 2002) and carries a recessive homozygous *pse-c* gene for resistance to group C (Hiehata et al., 2012). The extra resistance genes could potentially be identified through research on ‘Champagne’. Only ‘Champagne’ (*pse-c*/*pse-c*), when crossed with varieties susceptible to group C (*Pse-c*/*Pse-c*), exhibits resistance to group C (Hiehata et al., 2014). This resistance to group C was a quantitative trait (Hiehata et al., 2014), indicating that the resistance to group C is influenced by additional dominant genes in addition to the recessive gene *pse-c*. Pathogens are more likely to cause breakdowns against plant resistance genes for qualitative traits controlled by one or a few genes, through mutation. On the other hand, for plant resistance genes associated with quantitative traits, which are under the control of multiple genes, the possibility of breakdown occurring is less. Therefore, studying QTLs (Quantitative Trait Loci) is important for plant breeding (Pilet-Nayel et al., 2017).

Recently, single nucleotide polymorphisms (SNPs) have emerged as an invaluable tool in breeding programs for the identification of disease resistance genes (Laila et al., 2019). Technological advances have made the acquisition of SNP markers through Restriction-site Associated DNA Sequencing (RAD-Seq) both more efficient and cost-effective (Andrews et al., 2016). In our research group, SNP markers generated by RAD-Seq have been successfully employed to construct high-density linkage maps for the bronze loquat *Eriobotrya deflexa* (Fukuda et al., 2019) and to analyze the genetic diversity of loquat (Nagano et al., 2022). Additionally, in pear, a linkage map using SNP markers from RAD-Seq has facilitated QTL analysis of genes associated with sugar biosynthesis (Nishio et al., 2021).

This study aimed to identify the QTL regions associated with additional resistance genes that confer resistance to group C in ‘Champagne’. We conducted inoculation tests on individuals within the crossbred seedling population derived from ‘Tanaka’ (*Pse-c*/*Pse-c*) and ‘Champagne’ (*pse-c*/*pse-c*) as the cross parents. Additionally, the genotype of SNP markers in each individual was determined using the RAD-seq method, and linkage maps were constructed. Moreover, our QTL analysis identified novel gene regions conferring resistance to *Pseudomonas syringae* pv. *eriobotryae* group C.

## 2. Materials and methods

### 2.1. Plant material and DNA purification

One hundred twenty-three individuals from the cross of ‘Tanaka’ (*Pse-c*/*Pse-c*) × ‘Champagne’ (*pse-c*/*pse-c*) were utilized for genetic mapping. Genomic DNA was extracted from young leaves employing the cetyltrimethylammonium bromide (CTAB) method (Doyle and Doyle, 1987), followed by RNase treatment and phenol/chloroform extraction. The DNA concentration was determined using the Qubit dsDNA BR Assay Kit (Invitrogen, MA, USA) and adjusted to 20 ng/µl for subsequent library preparation.

### 2.2. Inoculation test and evaluation of resistance

The group C strain CG001 (Hiehata et al., 2002) was cultured on potato sucrose agar (PSA) medium at 25°C for one week. The bacteria were then suspended in sterile distilled water to a concentration of 10^8^ cfu/mL with 0.02% Tween 20 and needle-inoculated along the midrib on the abaxial side of three leaves per individual, targeting six sites per leaf. After inoculation, leaves were enclosed in vapor-deposited bags for approximately 24 hours to facilitate infection. Post-inoculation, the seedlings were maintained in a greenhouse to prevent other infections.

Resistance was evaluated two months after inoculation. The absence of dark brown canker disease symptoms was interpreted as resistance, while the presence of such symptoms indicated susceptibility. Resistance levels were scored as 0 (strong resistance) or 1 (weak resistance), based on the depth of the brown scab (Fig. 1A). Susceptibility levels were determined by the length of canker symptoms and categorized into three levels: 2 (up to 5.5 mm), 3 (5.6 mm to 7.5 mm), and 4 (more than 7.6 mm). Eighty-five individuals in 2021 and 123 in 2022 were inoculated to resistance evaluation.

**Fig. 1.**
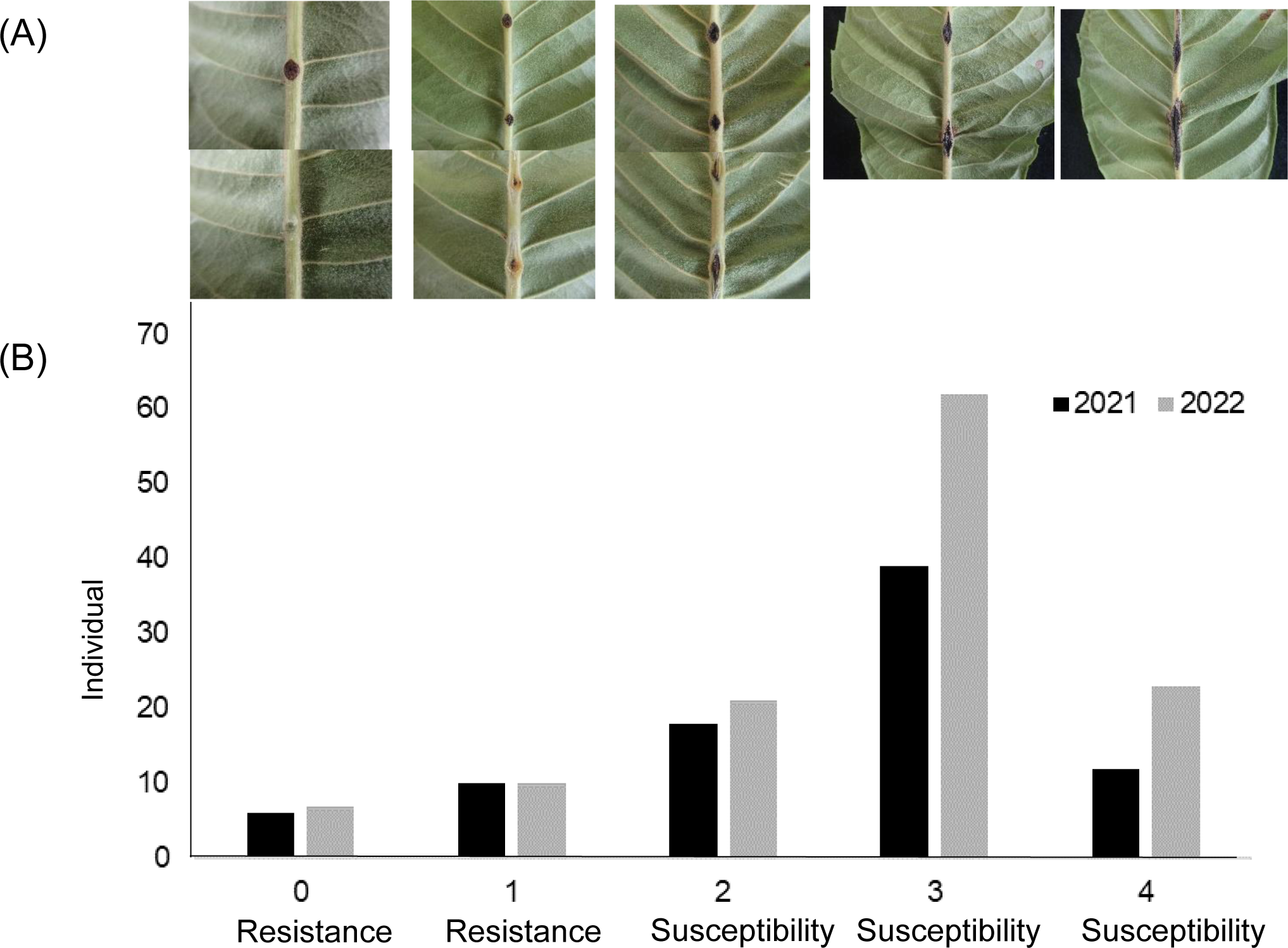
(A) Canker symptoms (disease index: 0 ∼ 4) at two months after inoculation with loquat canker. The absence of dark brown canker disease symptoms was interpreted as resistance 0 (strong resistance), 1 (weak resistance). Susceptibility levels were determined by the length of canker symptoms and categorized into three levels, 2 (up to 5.5 mm), 3 (5.6mm to 7.5 mm), 4 (more than 7.6 mm). (B) Frequency distribution of disease index with loquat canker. In the inoculation tests of 2021 (black) and 2022 (gray).

### 2.3. RAD-Seq analysis

The RAD-Seq library was prepared using the modified double-digest RAD-Seq method (Sakaguchi et al. 2015), which is an adaptation of the original methodology (Peterson et al. 2012). The sequencing was carried out by Macrogen (Seoul, Korea) on the HiSeqX platform (Illumina, San Diego, CA, USA), generating 151 bp paired-end sequences.

Initially, the sequencing data underwent quality control with the fastp tool (version 0.23.2), setting the sequence length option to 151 bp. After quality filtering, the sequences were aligned to the ‘Seventh Star’ reference genome (Jiang et al., 2020) using BWA software (version 0.7.17). The alignment files (in SAM format) were processed with Samtools (version 1.15.1) to sort them and convert them into BAM format.

For SNP detection and genotype analysis, the BAM files were processed using the “ref_map.pl” script from the Stacks software package (version 2.61) (Catchen et al., 2013). To accurately determine the genotype of each SNP marker across the population, the “populations” module of Stacks was utilized with specific parameters: a minimum allele frequency of 0.05 (--min-maf 0.05), a requirement for data presence in 80% of the population (-R 0.8), and outputs formatted for compatibility with JoinMap (--ordered-export, --map-type cp, and --map-format joinmap).

### 2.4. linkage mapping

JoinMap® 4.1 software (Kyazma B.V., the Netherlands; Van Ooijen, 2011) was employed for constructing the linkage map. SNP markers, observed in more than 50% of individuals and identified with the ‘nn×np’ and ‘lm×ll’ genotype, were analyzed using the cross-pollinated (CP) population option. With a LOD threshold of 10, markers were organized into 17 LGs. Linkage maps were then generated using the software’s default maximum likelihood mapping algorithm. The Kosambi mapping function was applied to convert recombination units into genetic distances. A sequence similarity comparison was conducted between the SNP markers on the linkage map and the ‘Seventh Star’ reference sequence (Jiang et al., 2020). Additionally, these sequences were compared to the apple draft genome (Zhang et al., 2019), which exhibits synteny with loquat, to align the linkage group numbers.

### 2.5. Statistical Analysis

To evaluate the reproducibility of the inoculation test, correlations between the disease indexes in 2021 and 2022 were calculated using Microsoft Excel spreadsheet software. The distribution’s normality of the disease index was assessed with the Shapiro-Wilk test, conducted using JMP Pro® 16.1.0 predictive analysis software (SAS Institute Japan Inc., Tokyo, Japan). Broad-sense heritability was estimated through an analysis of variance, which utilized variance components with genotypes and year as factors (Moriya et al., 2019). The formula:

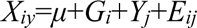

represents the phenotypic value for the *i*-th genotype in the *j*-th year, where *μ* is a constant, *G_i_* is the effect of the *i*-th genotype, *Y_j_* is the effect of the *j*-th year, and *E_ij_* is the error term for the *i*-th genotype in the *j*-th year, with *i* ranging from 1 to 123, and *j* from 1 to 2. If the genetic variance is *σ_g_^2^*, the annual variance is *σ_y_^2^*, and the error variance is *σ^2^*, then the environmental variance for the mean over *n* years, *σ_e_^2^* is calculated as (*σ_y_^2^*+*σ^2^*)/*n*, and the broad-sense heritability, *h_B_^2^*, is defined as *h_B_^2^*=*σ_g_^2^*/(*σ_g_^2^*+*σ_e_^2^*).

### 2.6. QTL Analysis

QTL analysis was conducted using MapQTL® 6.0 software (Van Ooijen and Kyazma, 2009). Due to the non-normal distribution of the evaluation traits, the Kruskal-Wallis (KW) test module was selected for initial analysis. For further evaluation of the detected QTLs, the restricted multiple QTL mapping (rMQM) module was employed to assess their contributions. The significance threshold for QTL identification was established at *P*=0.005, in accordance with the recommendation provided in the MapQTL manual for the KW test. A QTL was deemed significant if it was found in proximity to a DNA marker with a value below this threshold. The interval mapping (IM) method was used to ascertain the presence of QTLs within the 95% confidence interval of peaks, with LOD values exceeding the 95% threshold as determined by 1,000 permutation tests.

The contribution of QTLs to the 2-year average disease index in the ‘Tanaka’ × ‘Champagne’ population was assessed through one-way ANOVA, considering the genotype of the SNP marker closest to the identified QTL as a factor.

## 3. Results

### 3.1. Inoculation tests and resistance evaluation

We evaluated the resistance of 85 individuals in 2021 and 123 individuals in 2022. The results from the inoculation tests for these two years showed significant differences in disease resistance among the tested individuals (Fig. 1B). There was a strong positive correlation between the disease resistance observed in 2021 and 2022, with a correlation coefficient of 0.758. This indicates a consistent pattern of either resistance or susceptibility to the disease among the individuals over the two years.

The broad-sense heritability, calculated as 0.87 from the disease resistance data over the two years, points to a strong genetic influence on this trait. This high heritability suggests that the observed resistance has a substantial genetic basis.

Furthermore, the distribution of the disease resistance data from both years did not conform to a normal distribution, as evidenced by the Shapiro-Wilk test (*P*<0.05). This non-normal distribution indicates the complexity of the resistance trait being studied.

Namely, our hybridization experiment has shown that resistance to group C is inherited in a quantitative manner. This underscores the genetic diversity and complexity of resistance within the crossbred population.

### 3.2. SNP analysis by RAD-Seq

For the purpose of genetic mapping, we analyzed 123 individuals. Using double-digest RAD-Seq, we generated approximately 100 Gbp of data, averaging 814 Mbp per individual. The total number of reads was about 663 million, with an average of 5.4 million reads per individual. After processing the data with fastp and setting the read lengths to 151 bp, we retained 94 Gbp of high-quality sequence data, which averaged 764 Mbp per individual. The total number of reads after filtering was 622 million, with an average of 5.1 million reads per individual (Table Supplementary S1).

We mapped the filtered reads to the ‘Seventh Star’ reference sequence using the BWA tool. This mapping achieved an average coverage depth of 81.8×, with a maximum depth of 143.6× and a minimum of 31.6× (Supplementary Table S2). In total, we identified 4,409 SNP markers in loquat that were used for genetic mapping.

### 3.3. Linkage Mapping

We developed the genetic map of ‘Champagne’ using 1457 SNP markers. This map includes 1016 markers, covering a total length of 1300.9 centimorgan (cM) and achieving an average marker density of 1.4 cM (Fig. 2; Table 1). The lengths of the LGs varied, with the shortest being 48.1 cM (LG4) and the longest 107.4 cM (LG15). The number of markers mapped to each LG also ranged widely, from 32 markers on LG1 to 93 markers on LG3. Furthermore, the genetic map of ‘Tanaka’ contained 989 SNP markers. Sixteen LGs were identified, which covered a genetic distance of 1190.4 cM with an average marker density of 1.4 cM (Table 1: Supplementary Fig. 1)

**Fig. 2.**
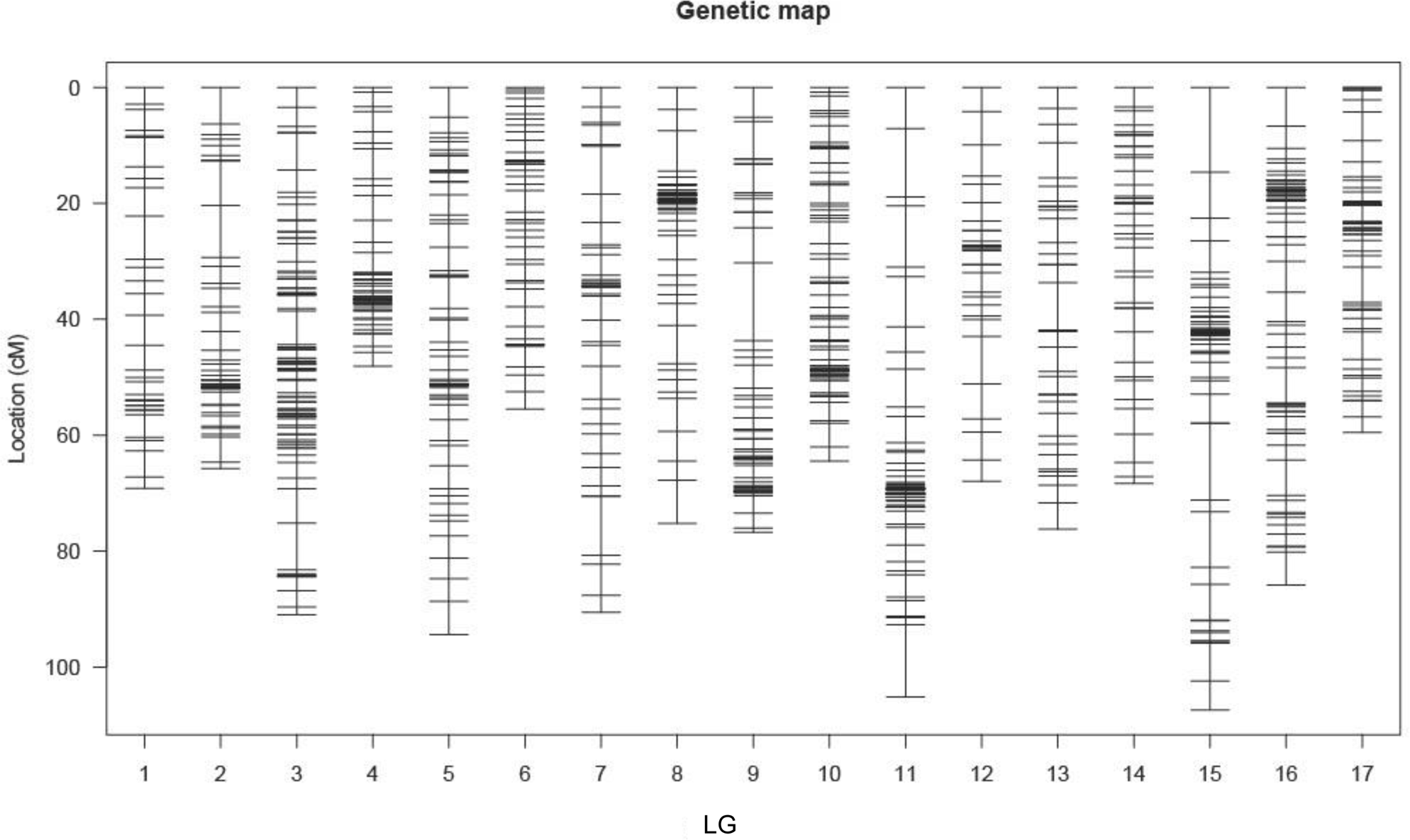
Genetic linkage map of ‘Champagne’. The vertical axis represents genetic distance (cM), and the horizontal axis represents linkage group numbers. Black lines indicate SNP markers.

**Table 1.** Summary of the linkage map of ‘Champagne’ and ‘Tanaka’.

|  |  | Linkage group |  |  |  |  |  |  |  |  |  |  |  |  |  |  |  |  | Total |
| --- | --- | --- | --- | --- | --- | --- | --- | --- | --- | --- | --- | --- | --- | --- | --- | --- | --- | --- | --- |
|  |  | LG1 | LG2 | LG3 | LG4 | LG5 | LG6 | LG7 | LG8 | LG9 | LG10 | LG11 | LG12 | LG13 | LG14 | LG15 | LG16 | LG17 |  |
| 'Champagne' | No. of mapped loci | 32 | 48 | 93 | 59 | 69 | 47 | 44 | 68 | 58 | 87 | 80 | 34 | 36 | 44 | 75 | 86 | 56 | 1016 |
|  | Length of linkage groups (cM) | 69.2 | 65.7 | 90.9 | 48.1 | 94.4 | 55.5 | 90.5 | 75.2 | 76.7 | 64.5 | 105 | 67.9 | 76.2 | 68.3 | 107.4 | 85.8 | 59.5 | 1300.9 |
|  | Marker density (cM/loci) | 2.2 | 1.4 | 1.0 | 0.8 | 1.4 | 1.2 | 2.1 | 1.1 | 1.3 | 0.7 | 1.3 | 2.0 | 2.1 | 1.6 | 1.4 | 1.0 | 1.1 | 1.4 |
| 'Tanaka' | No. of mapped loci | 106 | 75 | 16 | 47 | 111 | 64 | 49 |  | 68 | 64 | 60 | 56 | 75 | 82 | 47 | 17 | 52 | 989 |
|  | Length of linkage groups (cM) | 84.1 | 76.4 | 60.8 | 63.1 | 80.7 | 64.9 | 77.1 |  | 59.2 | 79.9 | 73.7 | 71.0 | 89.3 | 108.0 | 93.9 | 37.7 | 70.6 | 1190.4 |
|  | Marker density (cM/loci) | 0.8 | 1.0 | 3.8 | 1.3 | 0.7 | 1.0 | 1.6 |  | 0.9 | 1.2 | 1.2 | 1.3 | 1.2 | 1.3 | 2.0 | 2.2 | 1.4 | 1.4 |

### 3.4. QTL analysis

QTL analysis by the KW test in ‘Champagne’, revealed major QTLs in the same region of LG14 for both the 2021 and 2022 analyses (Fig. 3; Table 2; Supplementary Fig. 2). In the 2022 analysis and when averaging across the two years, the most notable QTL was found near the SNP marker SNPEjp207554, positioned 7.7 cM from the top of LG14 (Fig. 3; Table 2). In 2021, the highest K* value—a statistic indicating the strength of association between a marker and a trait—was near SNPEjp226440, located 16.8 cM from the top of LG14. The presence of QTLs in the upper region of LG14 was consistent across all datasets, with significance levels below *P*<0.005.

**Fig. 3.**
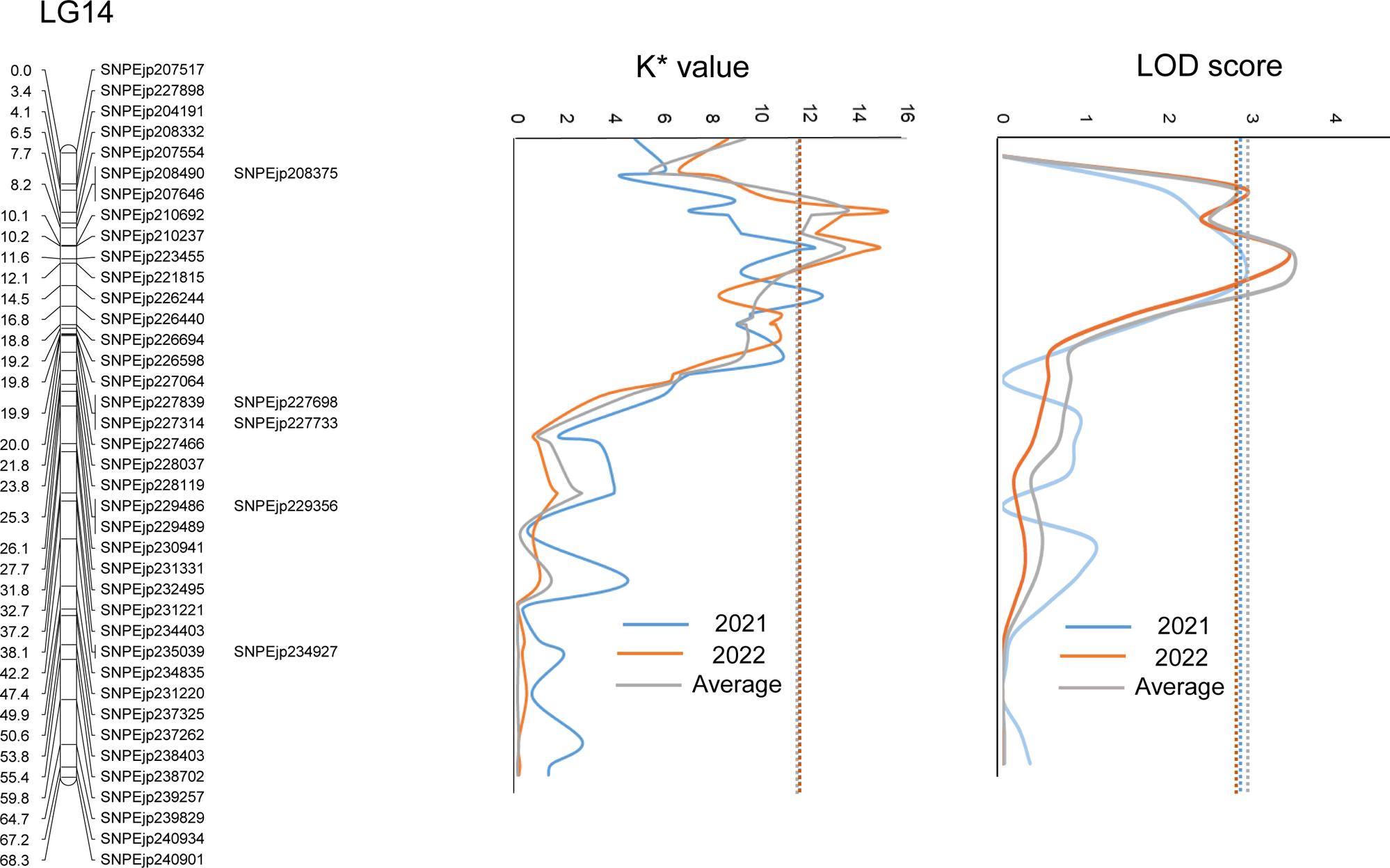
QTL for resistance against loquat canker identified in the genetic linkage group 14 in the Kruskal-Wallis test and interval mapping analyses. K values and LOD scores peaked on LG14 in 2021 (blue), 2022 (orange), and the two-year average (gray), confirming QTLs.

**Table 2.**
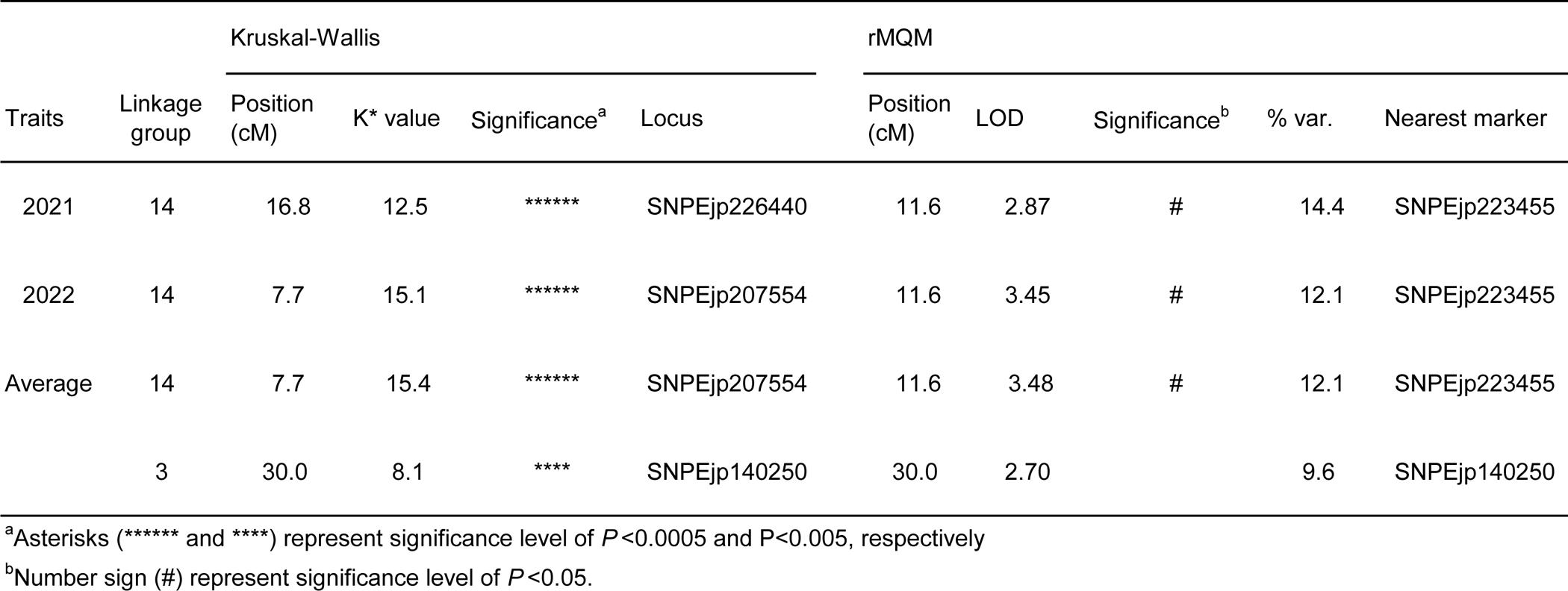
QTLs identified by the Kruskal-Wallis and rMQM methods for ‘Champagne’.

Further analysis using IM and rMQM methods confirmed the presence of QTLs with peak LOD scores surpassing the threshold in LG14 across three datasets. The SNP marker SNPEjp223455, located 11.6 cM from the top of LG14, showed the highest LOD score (Fig. 3; Table 2). This marker accounted for 6.9% of the variation in the disease index over a two-year average. We identified multiple minor potential QTLs across different linkage groups beyond LG14. Notably, the most significant of these was found in LG3, accounting for 3.7% of the variation in disease resistance over an average of two years (Table 2, Supplementary Fig. 2). In contrast, no stable QTL were observed in ‘Tanaka’.

## 4. Discussion

In this study, we identified new regions of resistance in the ‘Champagne’ cultivar against loquat canker, the most significant disease affecting loquat cultivation. This provides foundational knowledge for breeding resistant varieties. The resistance to group C studied this time is a quantitative trait, and as expected, we found that multiple genes were involved and the gene(s) with the most significant impact were situated on LG14.

*P. syringae*, a bacterium that infects various plants, causes significant diseases. In Arabidopsis, soybean, and tomato, qualitative resistance against *P. syringae* has been associated with specific genes known as *R* genes. These genes, including *RPM1* and *RPS2* in Arabidopsis, *Rpg1* in soybean, and *Pto* in tomato, feature nucleotide-binding sites (NB) and leucine-rich repeats (LRR), which are crucial for disease resistance (Mackey et al., 2002; Axtell and Staskawicz, 2003; Mackey et al., 2003; Ashfield et al., 1995; Selote and Kachroo, 2010; Martin et al., 1993). Additionally, research has shown quantitative resistance to *P. syringae* in other plants, such as kiwifruit, which implicates receptor-like serine/threonine protein kinases and other genes as potential resistance genes (Tahir et al., 2020). In tomato, while QTL regions have been identified using wild species, specific candidate genes have yet to be pinpointed (Thapa et al., 2015). The findings and related mechanisms from these studies on other plants may be relevant to the resistance of loquat against *P. syringae*.

Apples and loquats are part of the Malinae subtribe within the Maleae tribe, belonging to the Amygdaloideae subfamily of the Rosaceae family (Campbell et al., 2007; Potter et al., 2007; Liu et al., 2020). Reports have highlighted the presence of synteny, or conserved segments, between their genome sequences (Fukuda et al., 2019). The LG14 of the apple genome, which shares synteny with the linkage group where the identified QTL exists in this study, was discovered to contain a gene that provides resistance to powdery mildew caused by *Podosphaera leucotricha* (Calenge and Durel, 2006). Additionally, several resistance genes, including CC-NBS-LRR, NBS-LRR, NBS, and TIR-NBS-LRR, have been identified on LG14 of the ‘Golden Delicious’ apple genome (Perazzolli et al., 2014). This suggests that loquat might also carry similar resistance genes, potentially playing a role in defending against loquat canker.

By analyzing the above-mentioned results of other plants alongside the annotated loquat genome data (Jiang et al., 2020), especially within the newly identified resistance region on LG14, we can potentially uncover detailed mechanisms of resistance to *P. syringae*. Although pinpointing specific genes linked to qualitative traits presents challenges, previous successes in identifying genes for qualitative traits other than resistance to *P. syringae* in various plants provide optimism for future breakthroughs.

In related research on tomato (Thapa et al., 2015), genome comparisons across different strains of *P. syringae* revealed variations in effector proteins, which could be key to understanding infection and resistance mechanisms. For *P. syringae* pv. *eriobotryae*, which infects loquat, draft genomes for groups A, B, and C have been outlined (Tashiro et al., 2021). Investigating the unique effector proteins in the group C strain may provide further insights into the specific resistance mechanisms in loquat varieties.

The LOD scores for the newly detected QTL regions were relatively low, and confidence interval were broad. The power of QTL detection was influenced by several factors: the nature of the analyzed population, population size, the number and density of markers, missing data in RAD-Seq data, the number of genes involved in the QTL, the magnitude of the QTL effect, and environmental factors (Jiang et al., 2022). Notably, the QTL detection might have been affected by the low number of individuals in this study, for which resistance could be tested in 2021. However, a QTL was found in the upper 7.7 cM to 16.8 cM of LG14, marking an important discovery. Future studies should consider QTL-Seq utilizing the bulk method (Takagi et al., 2013) to further investigate the related genes.

In this study, we conducted a hybridization experiment with two cultivars, ‘Tanaka’ and ‘Champagne’, and constructed a linkage map of ‘Champagne’ consisting of 1016 markers. Previously, we developed a genetic linkage map of the bronze loquat (*Eriobotrya deflexa*) for a three-way cross of loquat (*E. japonica*) × (loquat × bronze loquat) (Fukuda et al., 2019). In the previous research, despite using RAD-Seq and simple sequence repeat (SSR) markers and analyzing genetically distant species, the linkage map consisted of 960 markers. The improvement in this study can be attributed to the use of 151 bp pair-end reads, compared to the 51 bp single-end reads used in the previous study.

‘Champagne’ shows resistance to multiple strains of *P. syringae* (Hiehata et al., 2002; Hiehata et al., 2012; Hiehata et al., 2014), making it an excellent source for resistance breeding. However, when grown in Japan, its fruit quality is inferior to that of other cultivars. Recently, ‘Harutayori’ was bred (Hiehata et al., 2016), exhibiting superior fruit quality to its ancestor ‘Champagne’. This cultivar is resistant to groups A, B, and C. It is expected to become widely adopted as an excellent source of resistance, especially in warm and humid cultivation environments where diseases are more likely to occur.

‘Champagne’ possesses the resistance genes *Pse-a* and *pse-c* against loquat canker, and these true resistance genes are being studied and utilized in resistance breeding. However, a limitation of these true resistance genes is that populations capable of overcoming the resistance emerge as the pathogen continuously evolves. To achieve stronger resistance, it is necessary to introduce QTLs that reduce disease infection rates in addition to resistance genes like *R* genes (Castro et al., 2003; Wulff et al., 2011). We believe the QTLs identified in this study will offer new avenues for resistance breeding against *P. syringae* pv. *eriobotryae*.

## CRediT author statement

**Shogo Koga:** Formal analysis, Writing - Review & Editing, **Ryusei Kawaguchi**: Formal analysis, **Tsunami Tanaka:** Formal analysis, **Shigeki Moriya:** Formal analysis, **Naofumi Hiehata:** Resources & Formal analysis, **Kozi Kabashima:** Formal analysis, **Atushi J. Nagano:** Formal analysis, **Yukio Nagano:** Formal analysis, Writing - Review & Editing, **Shinji Fukuda:** Formal analysis, Writing - Review & Editing, Project administration, Supervision

## Acknowledgements

This work was partially supported by the Matsushima Horticultural Development Foundation. This study was part of the dissertation submitted by the first author toward the partial fulfilment of their Ph.D. All authors provided consent for submission and publication.

## Data archiving statement

Sequences are available at the DDBJ Sequence Read Archive (https://ddbj.nig.ac.jp/resource/sra-submission/DRA015997)

---

**Supplementary Fig.1.** Genetic linkage map of ‘Tanaka’. The vertical axis represents genetic distance (cM), and the horizontal axis represents linkage group numbers. Black lines indicate SNP markers.

**Supplementary Fig.2.** QTL analyses for loquat canker disease index. Panel (A) indicate results obtained by Kruskal-Wallis analyses. Horizontal dotted lines indicate showing significant K*(*P*= 0.001). Panel (B) indicate results obtained by interval mapping analyses. Horizontal dotted lines indicate threshold scores (*P* = 0.05) obtained by permutation test.

