## Supplementary Fig. 1, 2, Supplementary Table S1, S2 for "Identification of new resistance QTL regions in loquat cultivar ‘Champagne’ against *Pseudomonas syringae* pv. *eriobotryae* group C"

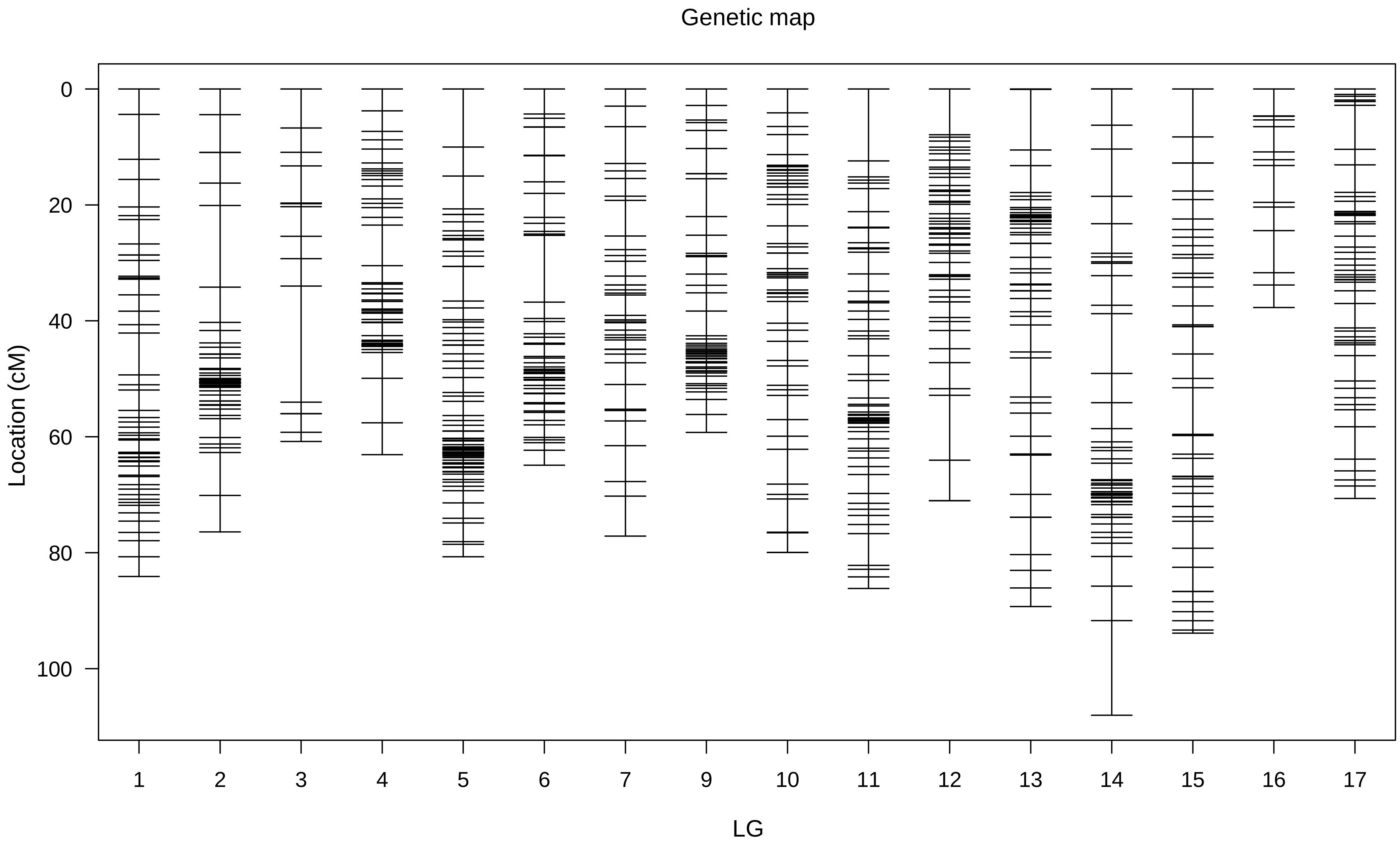

Supplementary Fig. 2.

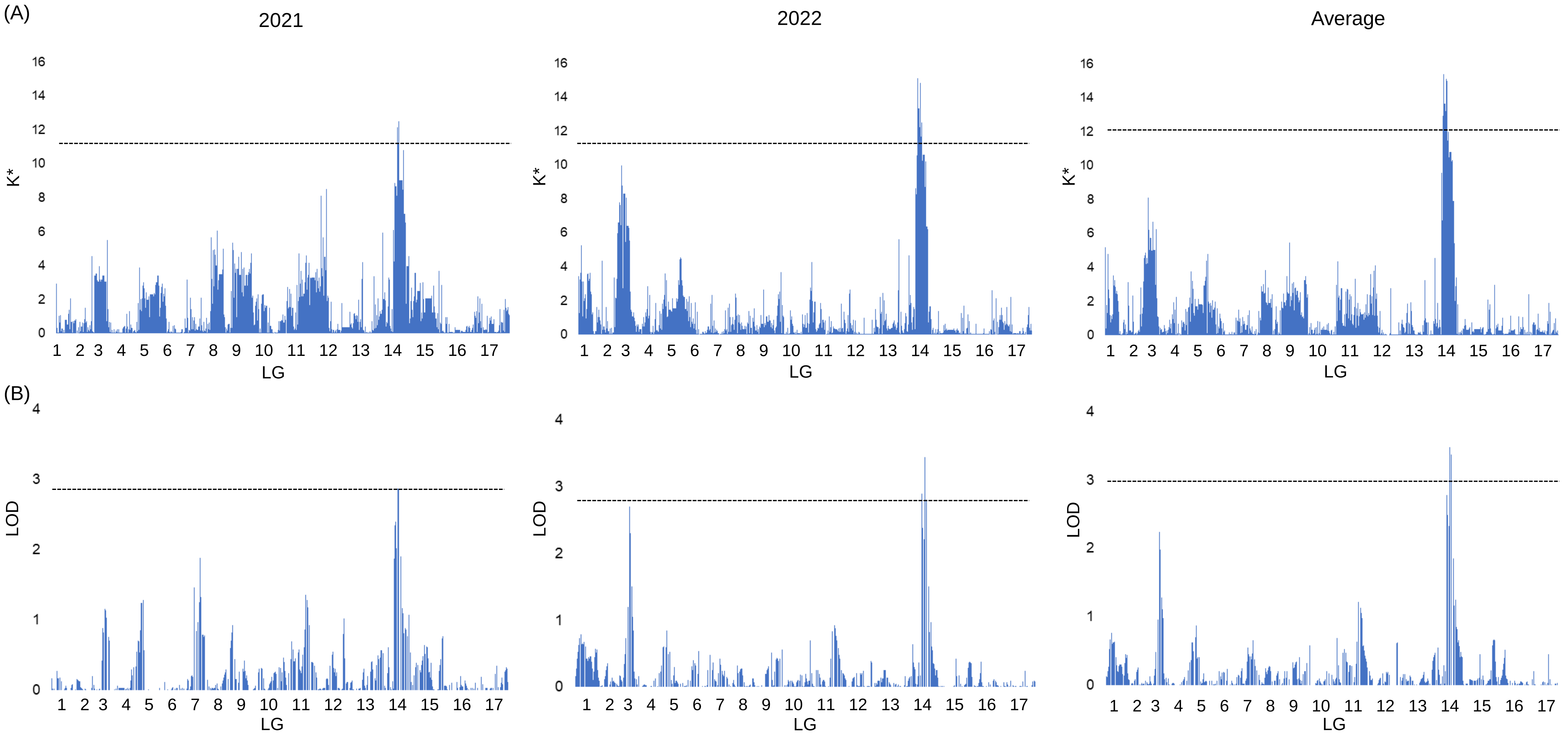

**Supplementary Table S1.** Number of filtered reads.

| Sample No. | Number of filtered reads |
|---|---|
| 1 | 5,536,836 |
| 3 | 4,909,477 |
| 8 | 5,922,361 |
| 10 | 3,794,022 |
| 11 | 4,836,065 |
| 12 | 5,039,783 |
| 15 | 3,424,415 |
| 16 | 3,409,399 |
| 17 | 4,693,981 |
| 18 | 5,121,937 |
| 19 | 4,774,951 |
| 20 | 7,368,431 |
| 21 | 6,747,614 |
| 22 | 5,139,057 |
| 23 | 5,593,691 |
| 25 | 3,566,056 |
| 26 | 3,891,979 |
| 27 | 7,897,890 |
| 28 | 9,660,407 |
| 29 | 5,046,082 |
| 30 | 3,878,939 |
| 31 | 5,006,304 |
| 32 | 5,196,350 |
| 33 | 2,879,261 |
| 34 | 3,779,070 |
| 35 | 6,360,212 |
| 36 | 6,853,371 |
| 37 | 6,512,749 |
| 38 | 6,490,977 |
| 39 | 6,258,207 |
| 40 | 4,786,198 |
| 41 | 5,385,118 |
| 42 | 6,056,673 |
| 45 | 6,087,284 |
| 48 | 4,945,108 |
| 49 | 5,161,939 |
| 50 | 5,048,611 |
| 51 | 4,373,614 |
| 52 | 3,982,707 |
| 53 | 3,801,837 |

| | |
|---|---|
| 54 | 2,968,766 |
| 55 | 4,559,768 |
| 56 | 5,069,479 |
| 57 | 4,974,943 |
| 58 | 4,862,188 |
| 59 | 3,989,551 |
| 60 | 4,628,839 |
| 61 | 5,168,210 |
| 62 | 4,586,906 |
| 63 | 4,209,156 |
| 64 | 4,154,840 |
| 65 | 4,095,722 |
| 66 | 4,122,560 |
| 67 | 4,357,178 |
| 68 | 4,282,206 |
| 69 | 3,995,442 |
| 70 | 4,581,833 |
| 71 | 4,605,199 |
| 72 | 4,431,969 |
| 73 | 5,082,187 |
| 74 | 4,811,582 |
| 75 | 4,402,244 |
| 76 | 4,150,906 |
| 77 | 3,501,752 |
| 79 | 5,097,284 |
| 80 | 4,094,617 |
| 81 | 4,156,025 |
| 82 | 4,533,234 |
| 83 | 3,610,079 |
| 85 | 4,964,573 |
| 92 | 3,506,551 |
| 94 | 3,891,371 |
| 99 | 6,572,751 |
| 100 | 7,306,051 |
| 102 | 5,411,872 |
| 103 | 5,710,035 |
| 107 | 3,467,254 |
| 108 | 3,904,657 |
| 109 | 5,301,356 |
| 110 | 5,699,896 |
| 111 | 5,957,993 |
| 113 | 4,653,134 |
| 115 | 3,913,517 |

| | |
|---|---|
| 116 | 4,731,890 |
| 117 | 5,747,569 |
| 118 | 4,787,316 |
| 119 | 3,401,815 |
| 120 | 4,053,364 |
| 121 | 4,083,601 |
| 122 | 4,420,105 |
| 123 | 6,067,753 |
| 124 | 6,208,824 |
| 125 | 3,858,598 |
| 126 | 4,317,038 |
| 127 | 5,692,544 |
| 128 | 4,951,200 |
| 129 | 4,505,732 |
| 130 | 3,687,297 |
| 131 | 3,957,501 |
| 132 | 5,038,434 |
| 133 | 4,915,924 |
| 134 | 4,679,955 |
| 138 | 4,597,925 |
| 139 | 5,502,342 |
| 140 | 5,345,846 |
| 141 | 5,707,532 |
| 142 | 6,798,093 |
| 144 | 4,341,829 |
| 146 | 3,921,546 |
| 147 | 6,171,996 |
| 149 | 8,007,025 |
| 150 | 6,519,040 |
| 151 | 5,847,004 |
| 152 | 5,981,927 |
| 153 | 5,071,906 |
| 154 | 3,087,303 |
| 155 | 3,911,160 |
| 156 | 5,706,029 |
| 157 | 7,222,581 |
| 158 | 8,102,154 |
| 159 | 8,277,045 |
| 160 | 8,704,329 |
| 161 | 7,876,824 |
| Total | 622,372,535 |
| Average reads | 5,059,939 |

**Supplementary Table S2.** Depth of coverage for each sample.

| Sample No. | Depth of coverage |
|---|---|
| 1 | 111.1 |
| 3 | 75.1 |
| 8 | 66.8 |
| 10 | 79.2 |
| 11 | 93.5 |
| 12 | 89.3 |
| 15 | 83.0 |
| 16 | 66.6 |
| 17 | 57.6 |
| 18 | 72.0 |
| 19 | 78.3 |
| 20 | 101.0 |
| 21 | 50.3 |
| 22 | 58.0 |
| 23 | 72.2 |
| 25 | 81.1 |
| 26 | 103.4 |
| 27 | 86.3 |
| 28 | 44.2 |
| 29 | 92.8 |
| 30 | 77.0 |
| 31 | 75.9 |
| 32 | 72.7 |
| 33 | 53.2 |
| 34 | 77.1 |
| 35 | 81.7 |
| 36 | 71.7 |
| 37 | 73.8 |
| 38 | 57.1 |
| 39 | 75.1 |
| 40 | 76.7 |
| 41 | 78.9 |
| 42 | 105.7 |
| 45 | 97.9 |
| 48 | 86.2 |
| 49 | 59.0 |
| 50 | 67.5 |
| 51 | 122.0 |
| 52 | 131.7 |
| 53 | 104.4 |

| | |
|---|---|
| 54 | 92.8 |
| 55 | 88.1 |
| 56 | 98.0 |
| 57 | 66.1 |
| 58 | 41.3 |
| 59 | 83.0 |
| 60 | 89.2 |
| 61 | 120.1 |
| 62 | 111.3 |
| 63 | 127.0 |
| 64 | 131.1 |
| 65 | 113.7 |
| 66 | 88.6 |
| 67 | 91.6 |
| 68 | 90.9 |
| 69 | 73.5 |
| 70 | 78.0 |
| 71 | 74.4 |
| 72 | 71.9 |
| 73 | 93.1 |
| 74 | 95.7 |
| 75 | 58.7 |
| 76 | 73.0 |
| 77 | 91.0 |
| 79 | 60.1 |
| 80 | 68.4 |
| 81 | 63.3 |
| 82 | 96.4 |
| 83 | 70.7 |
| 85 | 143.6 |
| 92 | 50.6 |
| 94 | 104.0 |
| 99 | 31.6 |
| 100 | 93.2 |
| 102 | 112.7 |
| 103 | 96.3 |
| 107 | 110.2 |
| 108 | 89.3 |
| 109 | 123.5 |
| 110 | 121.2 |
| 111 | 79.3 |
| 113 | 97.0 |
| 115 | 85.6 |

| | |
|---|---|
| 116 | 61.0 |
| 117 | 83.7 |
| 118 | 50.6 |
| 119 | 53.8 |
| 120 | 108.1 |
| 121 | 72.9 |
| 122 | 94.8 |
| 123 | 67.7 |
| 124 | 67.9 |
| 125 | 85.1 |
| 126 | 84.0 |
| 127 | 71.6 |
| 128 | 76.2 |
| 129 | 74.1 |
| 130 | 76.7 |
| 131 | 65.1 |
| 132 | 91.3 |
| 133 | 62.7 |
| 134 | 74.7 |
| 138 | 90.6 |
| 139 | 72.2 |
| 140 | 85.0 |
| 141 | 65.3 |
| 142 | 63.7 |
| 144 | 77.7 |
| 146 | 77.5 |
| 147 | 74.7 |
| 149 | 93.4 |
| 150 | 74.9 |
| 151 | 74.6 |
| 152 | 72.1 |
| 153 | 86.6 |
| 154 | 75.7 |
| 155 | 59.9 |
| 156 | 61.4 |
| 157 | 67.1 |
| 158 | 73.5 |
| 159 | 92.5 |
| 160 | 66.0 |
| 161 | 93.8 |
| Average reads | 81.8 |
